# *Saccharomyces cerevisiae* as a Platform for Assessing Sphingolipid Lipid Kinase Inhibitors

**DOI:** 10.1101/148858

**Authors:** Yugesh Kharel, Sayeh Agah, Tao Huang, Anna J. Mendelson, Oluwafunmilayo T. Eletu, Peter Barkey-Bircann, James Gesualdi, Jeffrey S. Smith, Webster L. Santos, Kevin R. Lynch

## Abstract

Successful medicinal chemistry campaigns to discover and optimize sphingosine kinase inhibitors require a robust assay for screening chemical libraries and for determining rank order potencies. Existing assays for these enzymes are laborious, expensive and/or low throughput. The toxicity of excessive levels of phosphorylated sphingoid bases for the budding yeast, *Saccharomyces cerevisiae*, affords an assay wherein inhibitors added to the culture media rescue growth in a dose-dependent fashion. Herein, we describe our adaptation of a simple, inexpensive, and high throughput assay for assessing inhibitors of sphingosine kinase types 1 and 2 as well as ceramide kinase and for testing enzymatic activity of sphingosine kinase type 2 mutants. The assay was validated using recombinant enzymes and generally agrees with rank order of potencies of existing inhibitors.

## Introduction

The sphingolipid kinases (sphingosine kinase (SphK) and ceramide kinase (CerK)) catalyze the transfer of phosphate from ATP to the primary hydroxyl of dihydrosphingosine (sphinganine), phytosphingosine, sphingosine, (SphK) or dihdyroceramides, N-acyl phytosphingosines and ceramides (CerK). Most eukaryotes have two sphingosine kinase genes while only a single ceramide kinase gene has been identified but some eukaryotes, e.g. *Saccharomyces cerevisiae*, lack detectable ceramide kinase activity. There has been recurrent speculation that these enzymes could be therapeutic targets, but their validation as such requires drug-like inhibitors, the discovery of which is dependent in part on facile biochemical assays.

A widely used assay for sphingolipid kinases involves incubating the enzyme with sphingosine and γ-[^32^P]ATP (or γ-[<^33^P]ATP), extracting non-polar compounds into an organic solvent mixture and detecting the radioactive product by autoradiography after normal phase thin layer chromatography [1]. In our experience this assay is both reliable and sensitive and it can be used with either recombinant enzymes or cell/tissue lysates. However, the assay is cumbersome, time consuming and too low throughput to support a medicinal chemistry campaign. Alternative assays have been developed to obviate the use of radioactive substrate and/or to eliminate the organic solvent extraction step. Such assays use fluorescently labeled [2] or biotinylated [3] substrates or capture the radiolabeled S1P on cation exchange paper [4], on streptadivin-coated paper [3] or on the walls of a 96 well plate [5]. Further, mass spectrometry based methods have been described [6], and a kit for detection of the ADP product are available (*e.g.* Promega’s ADP-Glo^TM^ Kinase Assay). All of these assays are more expensive and/or lower throughput relative to the yeast-based assay discussed herein.

Given our need for a higher throughput, inexpensive assay for sphingosine kinases to assess new chemical entities as potential inhibitors, our attention was drawn to a recent report by Kashem *et al*. [7] describing the use of the budding yeast, *Saccharomyces cerevisiae*, as a platform to screen for human SphK1 inhibitors. This assay takes advantage of the toxicity of excessive levels of phosphorylated LCBs (Long Chain Bases, *i.e.* sphinganine, phytosphingosine) to *S. cerevisiae* [8-11]. In wild type yeast, phospho-LCBs are degraded by two catabolic enzymes – a phosphatase (encoded by the *LCB3* gene) and a lyase (encoded by *DPL1*). Mutant yeast strains null for these gene products have high levels of phospho-LCBs and grow poorly [8,9] or not at all [11]. However, if the genes encoding the phospho-LCB generating kinases (*LCB4, LCB5*) are deleted also, normal growth is restored to the quadruple null mutant strain [8].

Forced expression of human SphK1 in the quadruple mutant (*LCB3*Δ *DPL1*Δ *LCB4*Δ *LCB5*Δ) strain results in severely retarded growth [7]. Importantly, Kashem *et al*. [7] demonstrated that growth of such a yeast strain can be rescued in a dose dependent fashion by adding an SphK1 inhibitor to the growth media. This result suggests that the products of the enzyme (phospho-LCBs), rather than forced expression of the human protein, are responsible for the toxicity. Herein, we report the extension of the findings of Kashem *et al*. to additional sphingolipid kinases, disclose uses of the assay beyond screening for SphK1 inhibitors and discuss the assay’s limitations and advantages based on our experience.

### Materials & Methods

The yeast strains used in this study include JS1256 (*MAT*a *his3Δ1 leu2Δ0 met15Δ0 ura3Δ0*) and CBY169 (*MATa leu2-3,112 ura3-52 his4 trp1 rme1 dpl1Δ::TRP1 lcb3Δ::LEU2 lcb4Δ::kanMX lcb5Δ::kanMX*) (the latter provided by Kyle Cunningham). The CBY169 strain was modified further by replacement of the *PDR5* gene with a selectable marker (confers resistance to clonat) to generate strain KYA1 (*MATa leu2-3,112 ura3-52 his4 trp1 rme1 dpl1Δ::TRP1 lcb3Δ::LEU2 lcb4Δ::kanMX lcb5Δ::kanMX pdr5Δ::natMX*).

The plasmids *pGAL-HsSPHK1*, *pGAL-HsSPHK2*, *pGAL-MmSphk1*, *pGAL-MmSphk1L277M*, *pGAL-MmSphk2*, *pGAL-HsCERK* and *pGAL-SK2_MOD* used in this study were constructed by sub-cloning DNA encoding the indicated translational open reading frames into the pYES2-FLAG-URA expression vector (provided by Dr. Cungui Mao). The encoded proteins all have an amino terminal FLAG epitope tag (DYKDDDDK) and their expression is under the control of the *GAL1,10* promoter. Excepting human and mouse SphK2, the DNA sequences are synthetic (from GeneWiz LLC (South Plainfield, NJ)) and were optimized for expression in *S. cerevisiae*. The Genbank accession number designations are as follows:

| lipid kinase | protein | nucleotide |
| --- | --- | --- |
| human SPHK1 | NP_001136074 | KY_859176 |
| human SPHK2 | NP_001191089 | NM_001204160 |
| mouse Sphk1 | NP_035581.1 | KY_859177 |
| mouse Sphk1 Leu277Met | none | KY_859178 |
| mouse Sphk2 | NP_001166032 | NM_001172561 |
| human CERK | NP_073603 | KY_859179 |
| human SK2_MOD | none | MF_642715 |

Yeast harboring plasmids were selected and maintained on synthetic complete media lacking uracil (SC-URA) with 2% glucose as the fermentable carbon source. After overnight growth at 30°C in this media, the cultures were diluted 1:100 into SC-URA media supplemented with 2% galactose and various concentrations of test inhibitor. After a further 24-48 hours incubation at 30°C, the extent of growth was assessed by measuring absorbance at 600 nm. Assays were conducted using either 3 – 5 mL of media in 16x150 mm glass tubes or 0.1 mL of media in Nunc U-bottom 96 well plates (VWR catalog # 62409-116) sealed with a gas-permeable membrane (Breathe Easy Plate Seals, VWR catalog # 102097-158). Tubes were incubated in a rotating drum while plates were agitated using a plate mixer. A_600_ for 96-well plate cultures was determined using a Molecular Devices SpectraMax M5 plate reader. Test compounds were dissolved in DMSO at 40-50 mM and diluted into water containing 0.1% fatty acid free bovine serum albumin. In all cases, the DMSO concentration in the assay was less than 2% (v/v).

### Construction of human SphK2 deletion mutants

To facilitate construction of deletion mutants of human SphK2, we altered the underlying DNA sequence to introduce a set of restriction enzyme sites that, on digestion and re-ligation, maintained the translational open reading frame (confirmed in each case by DNA sequencing). While some of the nucleotide changes were translationally silent, the altered human SphK2 (termed ‘SK2_MOD’) DNA encoded five amino acid changes (Glu71Asp, Gly138Ser, Glu146Asp, Glu437Gly, Gly623Ala).

### Determination of protein expression

The expression of plasmid-encoded sphingolipid kinases was determined using the protocol of Zhang *et al*. [12]. Yeast cells expressing deletion mutants of N-terminal FLAG-tagged human SphK2 were grown overnight on SC-URA media with 2% glucose. The cultures were diluted 1:100 into SC-URA media supplemented with 2% galactose in presence of an SphK2 inhibitor (0.5 μM SLM6031434) for 24-48 hours. Cells from 1 mL culture fluid were collected by centrifugation, re-suspended in 2 mL of 2M lithium acetate and incubated on ice for 5 minutes. This procedure was repeated and cells were suspended in 2 mL of 0.4M NaOH and incubated on ice for 5 minutes, re-centrifuged and the pellets were re-suspended in 0.25 mL of 1X Laemmli buffer and heated to 95°C for 5 minutes. After clarification by centrifugation, the supernatant fluid (10-15 μL) was loaded on 4-20% polyacrylamide gels and the resolved proteins were transferred onto a nitrocellulose membrane. Membranes were blocked with 5% (w/v) non-fat dried skimmed milk powder in TBS (Tris-buffered saline, pH 7.4) containing 0.1% Tween 20 for 1 hour at room temperature. After rinsing, membranes were incubated with monoclonal anti-FLAG M2-peroxidase antibody (Sigma-Aldrich #A8592) for an additional hour. After washing three times in TBS, the blot was developed by chemiluminesence using a commercial kit (PerkinElmer Western Lightning).

### Quantification of yeast sphingolipids by liquid chromatography / mass spectrometry (LCMS)

Cell pellets (approximately 5 million cells) were mixed with 2 mL of a 3:1 methanol/chloroform mixture and transferred to a capped glass vial. To this suspension was added 10 μL of internal standard solution containing 1 μM d7S1P (deuterated sphingosine 1-phosphate) (Avanti Polar Lipids). The mixture was homogenized in a bath sonicator for 10 min and incubated at 48°C for 16 h. The mixture was then cooled to room temperature and mixed with 200 μL of 1 M KOH in methanol. The samples were again sonicated and incubated at 37°C for 2 h. After this time, the samples were neutralized through the addition of 20 μL of glacial acetic acid and transferred to 2 mL microcentrifuge tubes. Samples were then centrifuged at 10,000 x g for 10 min at 4°C. The pellets were discarded, the supernatant fluid was collected in a separate glass vial and the solvent was evaporated under a stream of nitrogen gas. Immediately before LCMS analysis, the material was dissolved in 300 μL of methanol, clarified by centrifuged at 12,000 x g for 12 min at 4°C, and analyzed by LCMS.

Analyses were performed using a triple quadrupole mass spectrometer (Sciex 4000 Q-Trap) coupled to a Shimadzu LC-20AD liquid chromatography system. A binary solvent gradient with a flow rate of 1 mL/min was used to separate sphingolipids by reverse-phase chromatography using a Supelco Discovery C18 column (50 mm × 2.1 mm, 5 μm bead size). Mobile phase A consisted of water/methanol/formic acid (79.9:20:0.1, by volume), whereas mobile phase B was methanol/formic acid (99.9:0.1, v/v). The run started with 100% solvent A for 0.5 min after which solvent B was increased linearly to 100 % solvent B over 5.1 min and held at 100 % solvent B for 4.3 min. The column was finally re-equilibrated to 100% solvent A for 1 min. Sphingolipids were detected using MRM (multiple reaction monitoring) methods as follows: d7S1P (387.4→271.3); dhS1P (382.4→266.4); dhSph (dihydroSphingosine) (302.5→60.0); phytoS1P (398.3→300.4) and phytosSph (318.4→282.3).

## Results

In adopting this assay platform, we first attempted to replicate the results of Kashem *et al*. [7] using both human SphK1 and SphK2. We were successful, specifically, we observed that forced expression (by growth in media wherein the sole fermentable carbon source is galactose) of either human isoform in the quadruple null strain (CBY169) inhibited growth, while forced expression of a sphingosine kinase was not severely growth inhibitory so long as either LCB-phosphate catabolic gene (*DPL1* or *LCB3*) was intact (Fig. 1). We extended this observation to another sphingolipid kinase, human ceramide kinase (HsCerK), the expression of which was growth inhibitory to our standard laboratory strain, JS1256 (‘WT’).

**Fig 1:**
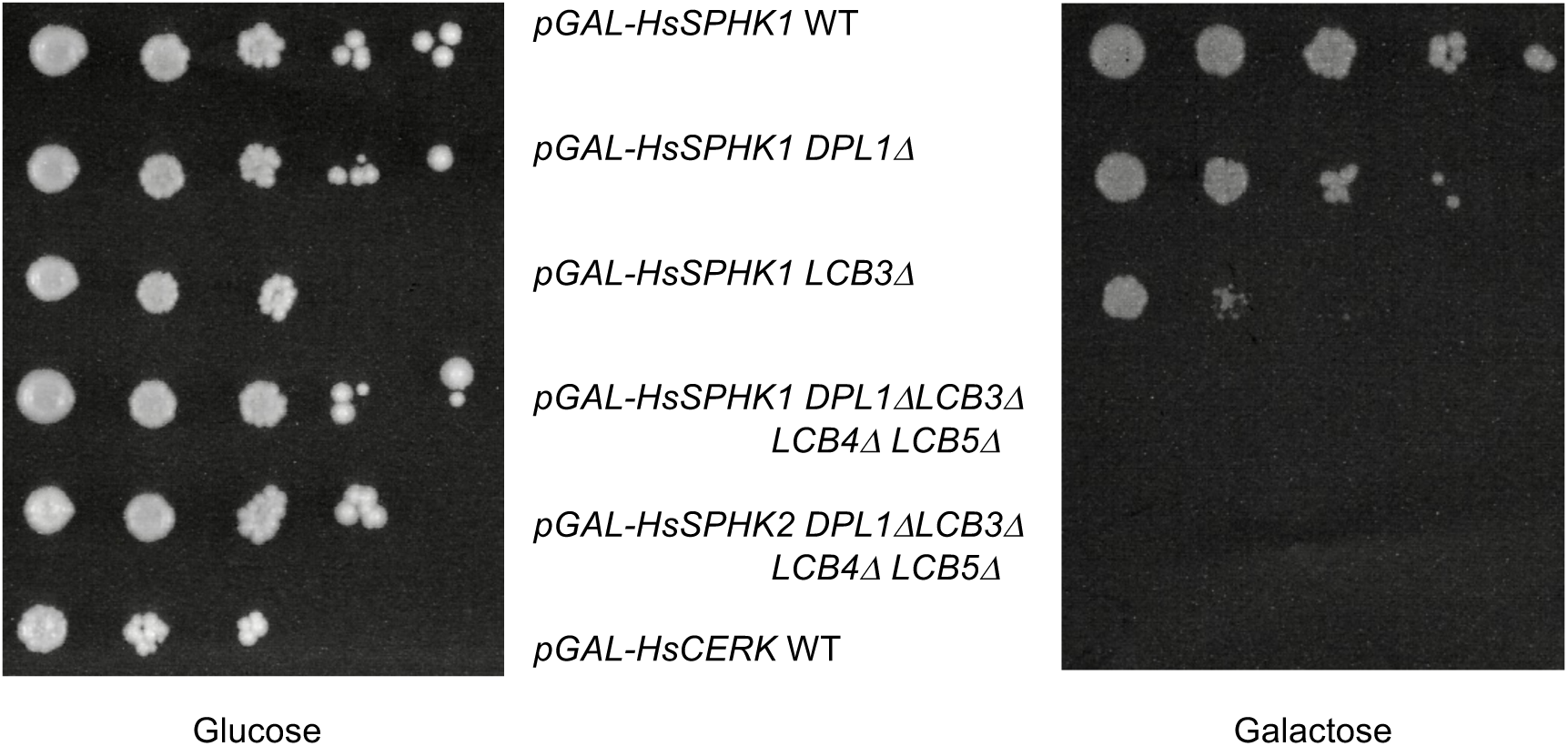
Growth of yeast strains on glucose or galactose SC media. Cultures were grown in SC-URA+GLU media to saturation, serially (10-fold) diluted in water and 2.5 μL was applied to solid SC-URA media containing GLU or GAL.

We then tested three widely used SphK inhibitors: PF-543 (SphK1 selective), SKI-II (non-selective) and ABC294640 (SphK2 selective) as well as Genzyme compound 9ab (SphK1 selective) (see Table 1 for structures and attendant literature citations) using the CBY169 strain transformed with either human SphK1 or SphK2 encoding plasmids. Among this set of four inhibitors, only compound 9ab restored full growth of the human SphK1 expressing strain in galactose media (Fig. 2A) with an EC_50_ value of 0.9 μM, which is similar to the K_I_ value (1.4 – 1.7 μM) that we determined previously for this compound using recombinant human SphK1 [5].

**Table 1:**
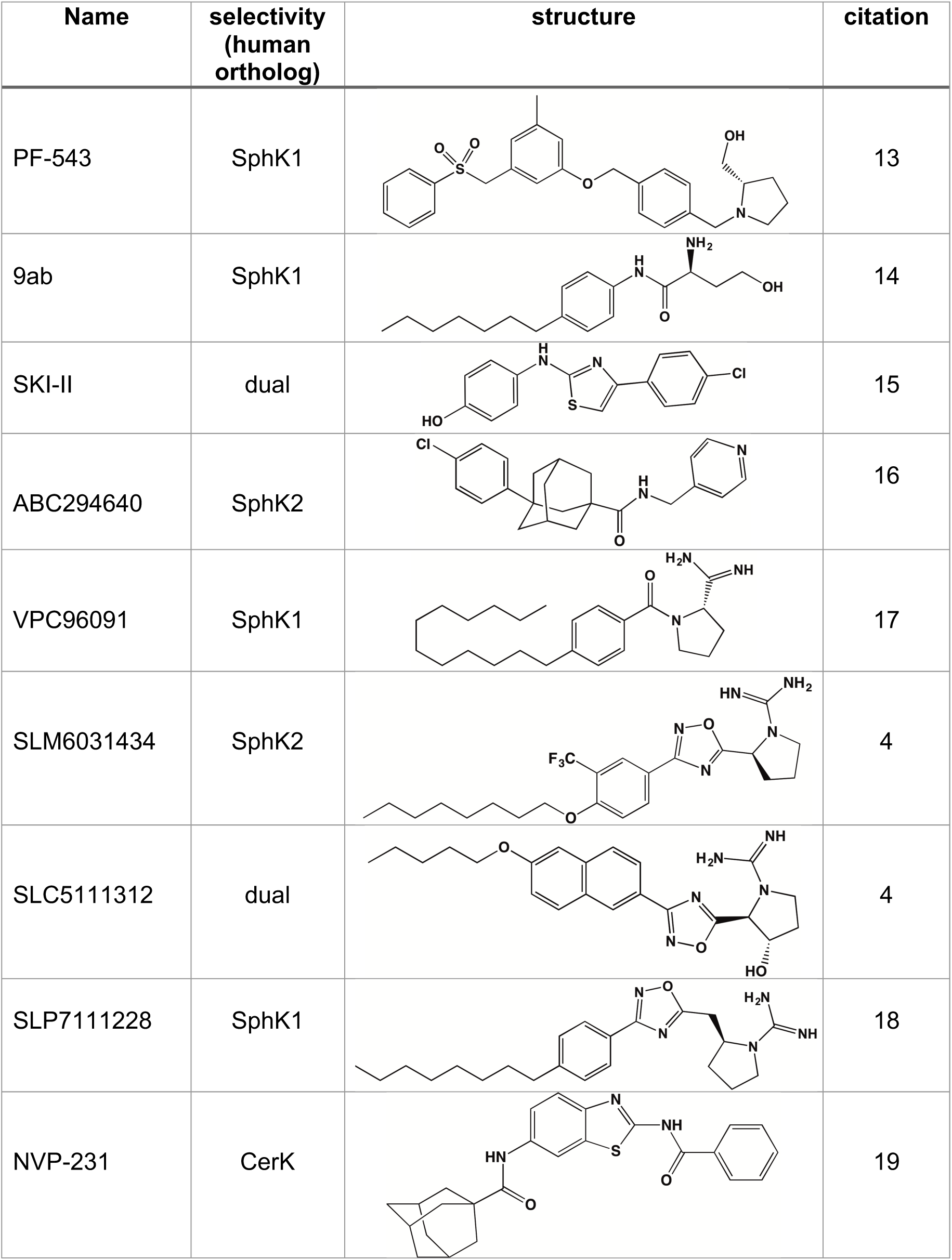
Sphingolipid kinase inhibitors used in this study

| Name | selectivity<br>(human<br>ortholog) | structure | citation |
| --- | --- | --- | --- |
| PF-543 | SphK1 |  | 13 |
| 9ab | SphK1 |  | 14 |
| SKI-II | dual |  | 15 |
| ABC294640 | SphK2 |  | 16 |
| VPC96091 | SphK1 |  | 17 |
| SLM6031434 | SphK2 |  | 4 |
| SLC5111312 | dual |  | 4 |
| SLP7111228 | SphK1 |  | 18 |
| NVP-231 | CerK |  | 19 |

The failure of PF-543, SKI-II and ABC294640 to rescue growth might be due to either cytotoxicity or failure of the compounds to accumulate (penetration – (extrusion + metabolism)) sufficiently in the yeast cells. To assess toxicity, we cultured the CBY169 strain without SphK plasmids in galactose media with these three compounds at concentrations up to 100 μM. In this experiment, we found only SKI-II to be toxic. Specifically, when present at concentrations above 3 μM, SKI-II inhibited growth (not shown), which obviates the use of this assay for assessing this low potency (K_I_ 12 - 30 μM [5]) compound. In an attempt to increase accumulation of compounds in the yeast cells, we modified CBY169 to generate a new strain (KYA1) wherein *PDR5* was deleted. PDR5p is an ABC (ATP Binding Cassette) transporter that confers resistance to a variety of xenobiotics [20]. This gene is commonly deleted in *S. cerevisiae* to decrease extrusion of test compounds. As shown in Fig. 2B, PF-543 restored full growth of the KYA1 strain expressing human SphK1 in a dose dependent manner, albeit with a EC_50_ value (5.7 μM) that is about three log orders higher than the reported K_I_ value of this compound [12]. ABC294640, although also not cytotoxic, failed to rescue growth of KYA1 yeast expressing HsSphK2 at concentrations up to 100 μM (not shown).

**Fig 2:**
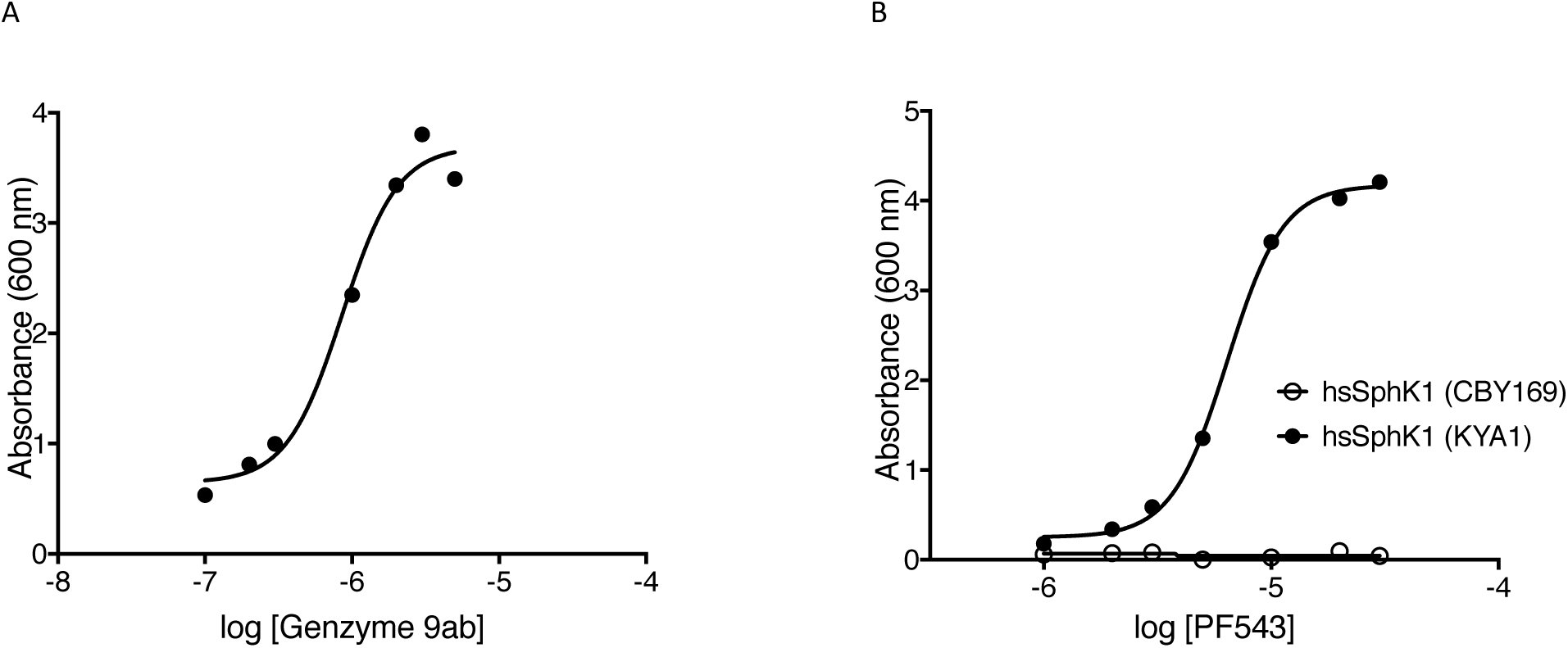
A - Rescue of growth of strain CBY169 expressing human SphK1 by 9ab; B-rescue of growth of strain KYA1, but not CBY169, expressing human SphK1 by PF-543

To investigate the yeast assay further, we tested additional inhibitors including VPC96091, SLP7111228 and SLC5111312 on human SphK1 expressing yeast. As documented in Fig. 3A, each of these compounds restored full growth in a dose dependent mannder and the rank order potency of the compounds was the same as determined at recombinant human SphK1 (SLP7111228 > VPC96091 > SLC5111312) [4,16,17]. In addition, we tested SLC5111312 and SLM6031434 at human SphK2 and found both compounds likewise rescued growth (Fig. 3B) with the same rank order potency as established with K_I_ values (SLM6031434 > SLC5111312). Unlike PF-543, the potency of SLM6031434 was the same regardless of PDR5p status (CBY169 vs. KYA1 strains) (Fig 3C).

We next asked whether ceramide kinase (CerK) expression effects the growth of *S. cerevisiae* and, if so, whether this phenotype could be reversed by adding a CerK inhibitor to the media. To our knowledge, neither ceramide kinase activity nor the predicted products of the enzyme (dihydroceramide 1-phosphate, phytoceramide 1-phosphate) have been observed in *S. cerevisiae*. When we forced expression of human CerK in our standard laboratory strain (JS1256) by culturing *pGAL-HsCERK* harboring yeast in galactose media, the cultures failed to grow (see Fig. 1) but growth was restored in a dose dependent fashion by addition of the CerK inhibitor, NVP-231 [19] to the culture media (Fig 3D). This result suggests that human CerK expression is toxic to yeast due to dihydroceramide 1-P and/or phytoceramide 1-P accumulation rather than toxicity of the human protein. We note that a problem intrinsic to ceramide biochemistry, *i.e.* the difficulty in manipulating naturally occurring ceramides, which are water-insoluble, is circumvented with the yeast assay.

**Fig 3.**
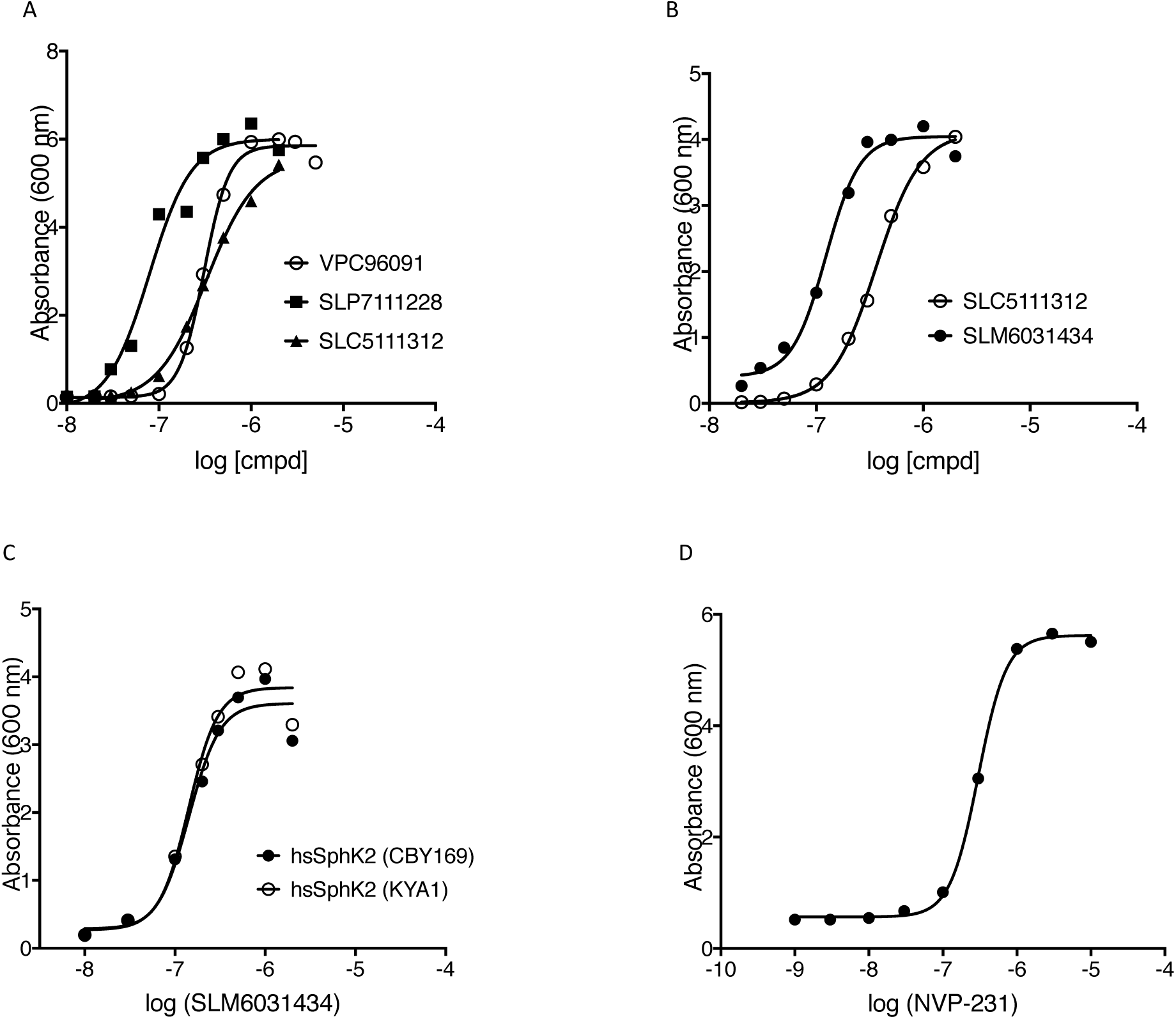
A - rescue of human SphK1-expressing yeast, B, C – rescue of human SphK2-expressing yeast and D – rescue of CerK-expressing yeast; C compares CBY169 and KYA1 strains; all other strains were CBY169

The data presented in Figures 2 and 3 are from single point assays, that is, only one determination was made at each inhibitor concentration. To estimate the variability of the assay in our hands, we performed 4 independent assays wherein the SphK2 selective inhibitor, SLM6031434, was used to rescue growth of human SphK2-expressing CBY169 strain yeast and compared the EC_50_ values so determined. These values were (in nM) 78, 127, 142, 144 (mean = 123, 95% confidence interval 92-153 nM). A second estimation of variability was achieved by performing the assay in 96 well plates whereby the SphK1 inhibitor, VPC96091, was assayed in triplicate. The variation among the data points is shown in Fig. 4. We note that the yeast cultures in the plates achieve nearly the same density as in individual 16 x 150 mm tubes, but this is not revealed in the maximum absorbance measurements from plate readings because of the limited linear range of the plate reader (culture media from tube assays was diluted 10-fold prior to determining A_600_).

To confirm that forced expression of a sphingosine kinase resulted in high levels of phosphorylated LCBs, we cultured yeast harboring either a *pGAL-HsSPHK1* or a *pGAL-HsSPHK2* plasmid in glucose or galactose media and measured cellular sphingoid bases and their phosphorylated analogs by LCMS. As documented in Figure 5, phospho-LCBs accumulated in galactose media, and their levels were decreased by inclusion of an inhibitor in the culture media (SphK1 inhibitor (VPC96091); SphK2 inhibitor (SM6031434)).

**Fig 4:**
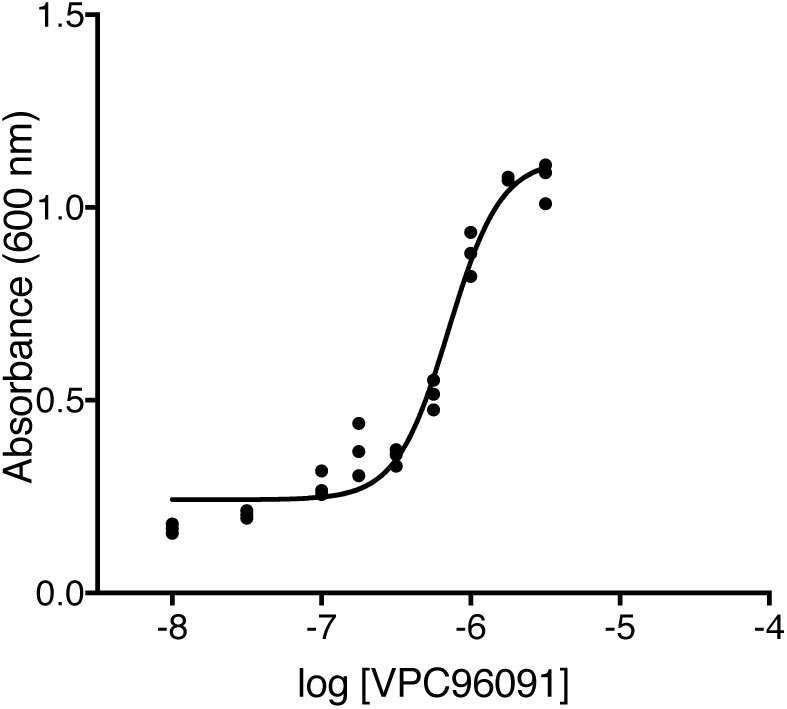
plate assay using VPC96091 and human SphK1

**Figure 5:**
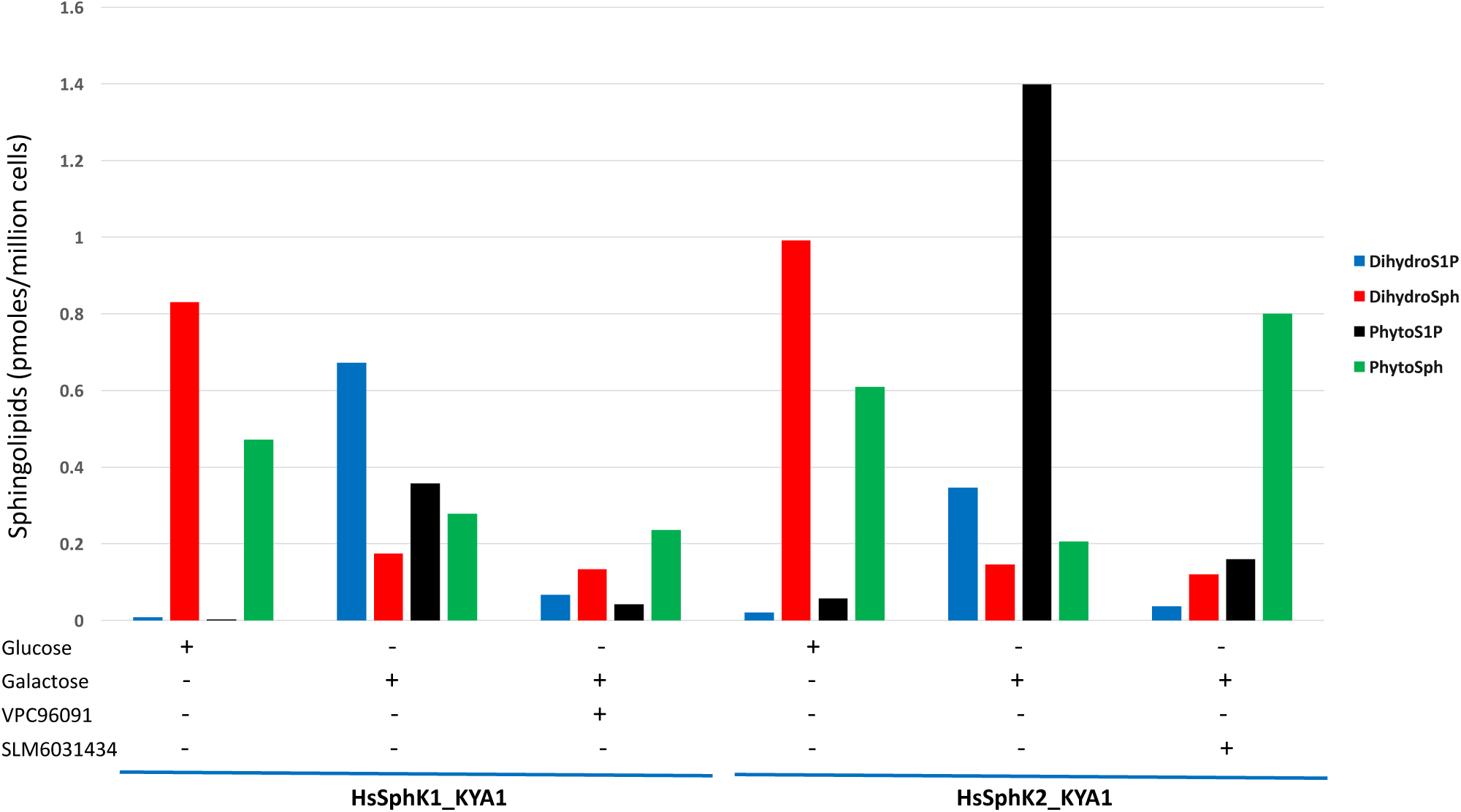
Yeast were cultured for 6 h in the indicated media; inhibitors were present at 300 nM. Sphingolipids in cell pellets were analyzed by LCMS (see Methods for details).

Encouraged by our success with *S. cerevisiae* as a sphingolipid kinase inhibitor assessment tool, we considered further applications of the yeast system for investigating sphingolipid kinase biology. The yeast system enables rapid interrogation of mutant enzymes for activity (*i.e.* inhibition of growth on galactose) and the potency of inhibitors at mutant enzymes (*i.e.* rescue of growth by inhibitors). We used this property of the system to investigate two questions: First, what amino acid(s) contribute to species difference in the potency of SphK1 inhibitors, specifically the lower potency of some SphK1 inhibitors at mouse (*vs.* human) SphK1? Second, what is the minimum size of a functional SphK2 enzyme, specifically, could we generate a functional SphK2 (72 kDa) that is the same size as SphK1 (48 kDa)?

We have observed that while SLC5111312 is equipotent at human and rat SphK1 and SphK2, it is a 20-fold SphK2 selective inhibitor in the mouse due to decreased potency at mouse SphK1 [4]. Human and rodent SphK1 amino acid sequences are about 80% identical while mouse and rat SphK1 are 90% identical. Alignment of the human, rat and mouse SphK1 amino acid sequences (not shown) reveals that position 277 is leucine in mouse but methionine in human and rat. The crystal structures of human SphK1 [21,22] indicate that this amino acid contributes to the sphingosine binding site in the enzyme (SLC5111312 is competitive with sphingosine). To test the hypothesis that Leu277 is responsible for lower potency of SLC5111312 at mouse SphK1, we obtained the mouse SphK1 L277M mutant, determined that its expression was, as expected, growth inhibitory for the CYA169 and KYA1 strains, and determined the potency of two inhibitors at the mutant and WT mouse enzymes. As depicted in Fig. 6A, VPC96091, which is an SphK1 inhibitor that does not discriminate between the mouse and human orthologs, is equipotent at wild type and mutant (L277M) mouse SphK1 while SLC5111312 is distinctly more potent at the mutant mouse SphK1 (6B). The EC_50_ values for SLC5111312 were similar for the mutant mouse enzyme and the human enzyme (see Fig. 3B).

**Fig. 6.**
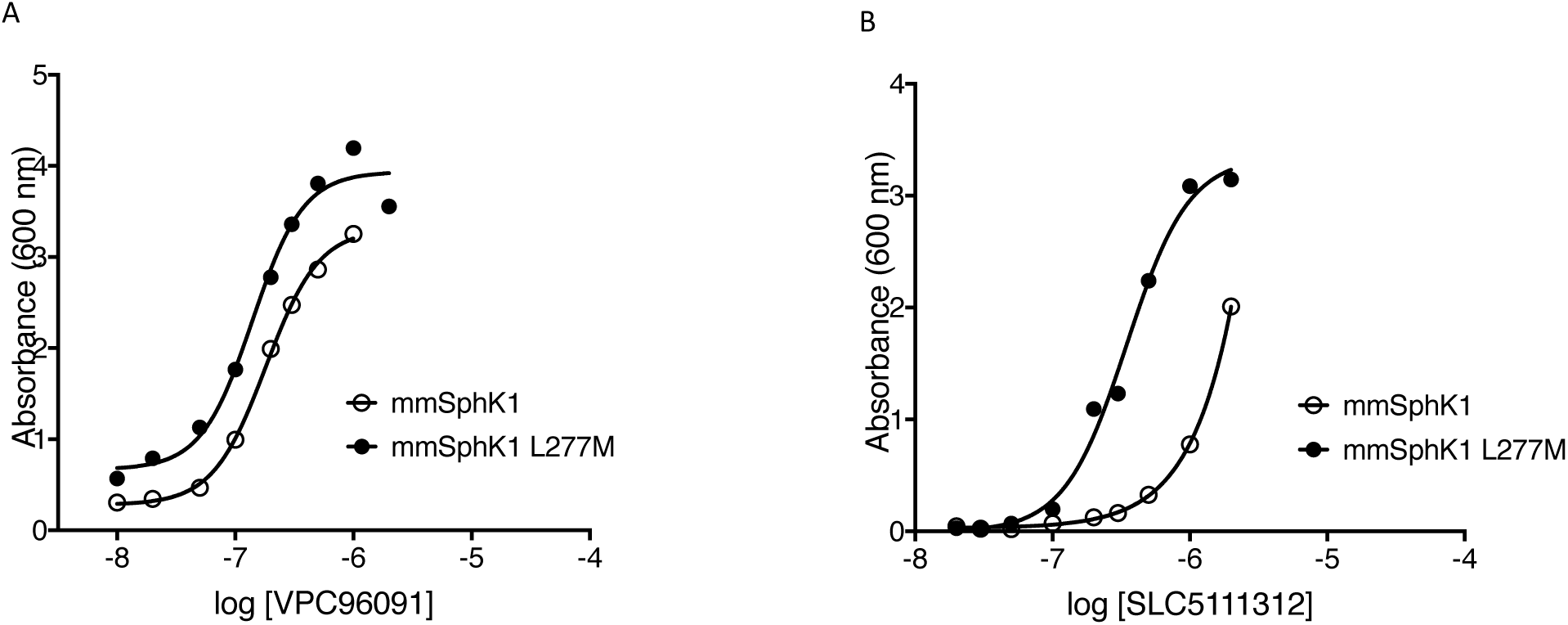
Rescue of mouse SphK1 and mouse SphK1L277M expressing yeast with VPC96091 (A) or SLC5111312 (B).

Human SphK2 is considerably larger than human SphK1 due to an extended (about 130aa) N-terminus and a centrally located 135aa segment. Because both enzymes catalyze the same reactions with similar kinetic constants [23], we presumed that deletion of much of the additional amino acid residues in SphK2 would not eliminate enzyme activity. However, we were less confident in predicting whether growth inhibition driven by smaller SphK2 mutants would be rescued by SphK2 inhibitors. To estimate the minimal size of functional human SphK2, we modified the encoding DNA to generate a set of mutants lacking portions of the non-SphK1 overlapping amino acids of human SphK2. The deleted regions are illustrated in Fig. 7A.

The expression of each deletion mutant was confirmed by western blotting (Fig. 7B). Each deletion mutant was tested in the yeast assay for activity (inhibition of growth on galactose media) and, if active, for rescue by the SphK2-selective inhibitor, SLM6031434. The concentration activity curves from the rescue experiment are presented as Fig. 8, and the results of the deletion study are summarized in Table 2.

**Fig. 7A.**
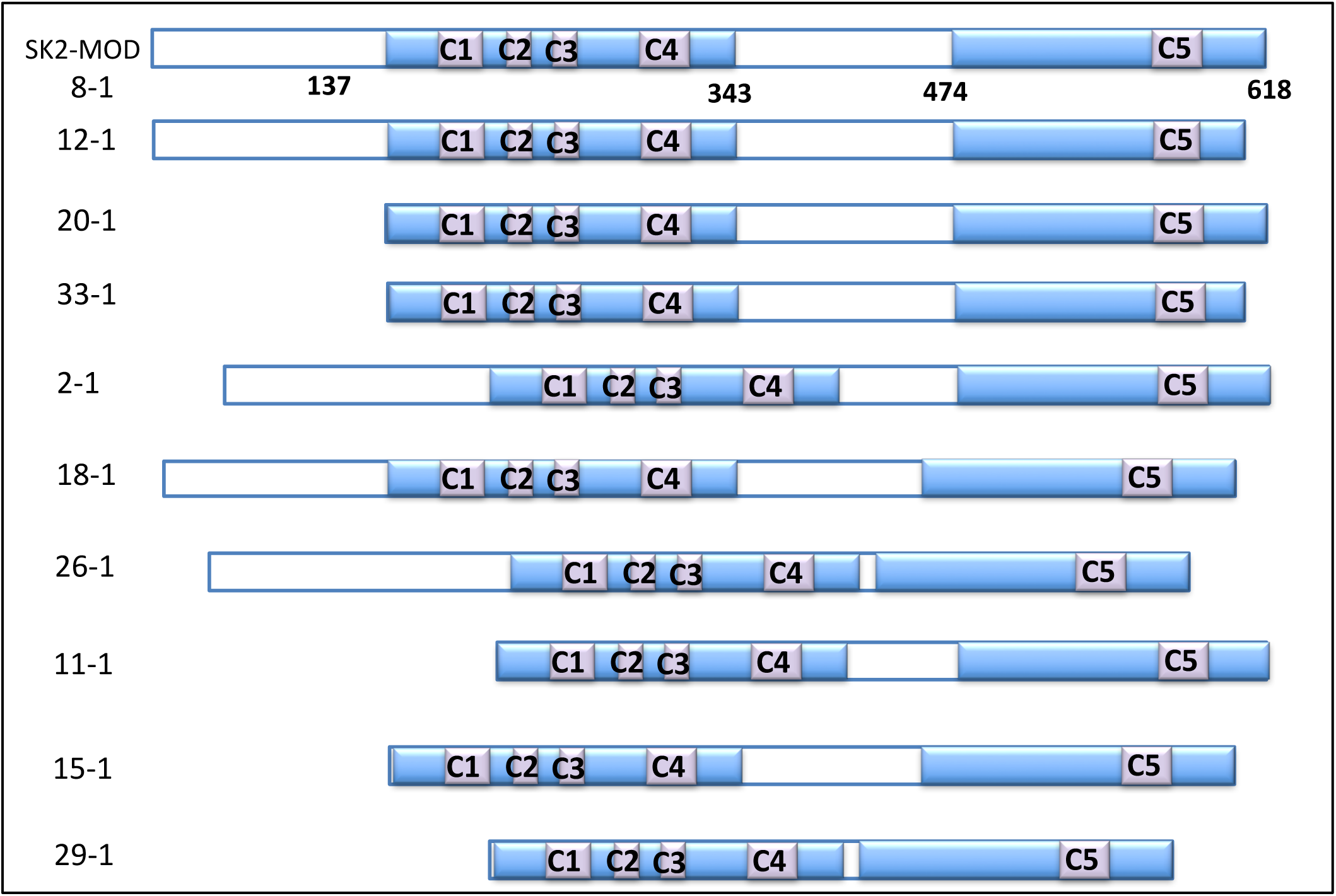
Diagram of human SphK2 deletion mutants; 8-1 represents SphK2_MOD. Shading indicates regions are common to human SphK1 and human SphK2 (ca. 50% identical aa) while open bars indicate sequence unique to SphK2. C1-C5 indicate amino acid stretches that are conserved among sphingosine kinases across multiple vertebrate species [23].

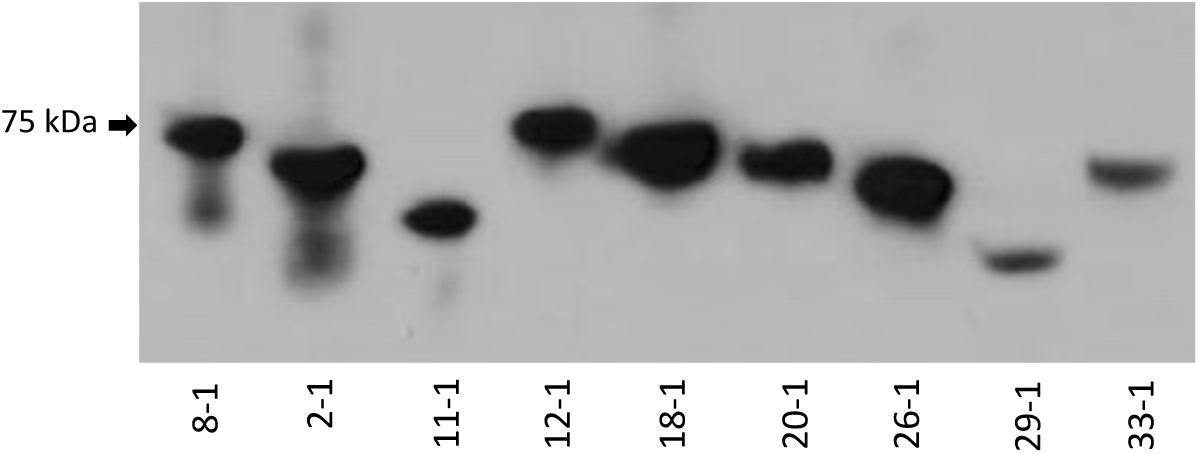
7B –Expression of deletion mutants: Yeast were grown to saturation in galactose media supplemented with 0.5 μM SLM6031434 except 33-1, which grew in galactose media without inhibitor. The volume of yeast extract (see Methods) loaded onto the gel was as follows: 8-1, 12-1, 18-1 26.1, 0.002 mL; 11-1 0.003 mL; 2-1, 20-1 0.004 mL; 29-1 0.02 mL; 22-1 0.01 mL. Note that growth of deletion mutant 15-1 was not rescued by SLM6031434.

**Table 2.** Activity of human SphK2 deletion mutants

| Mutant designation | Deleted segment (aa) | growth on galactose | Rescue by SLM6031434 (EC<sub>50</sub> (μM)) | Expression (FLAG epitope) |
| --- | --- | --- | --- | --- |
| - | full length (wild type hSK2) | no | 0.14 | + |
| 8-1 | full length (hSK2_MOD) | no | 0.10 | + |
| 20-1 | 12 - 137** | no | 0.05 | + |
| 2-1 | 371 - 436 | no | 0.28 | + |
| 18-1 | 437 - 481 | no | 0.10 | + |
| 26-1 | 371 - 481 | no | > 1 | + |
| 12-1 | 623 - ter | no | 0.10 | + |
| 11-1 | 12 - 137 & 371 - 436 | no | 0.28 | + |
| 15-1 | 12 - 137 & 437 - 481 | no | no | n.d. |
| 29-1 | 12 - 137 & 371 - 481 | no | > 1 | + |
| 33-1 | 12 - 137 & 623 - ter | yes | - | + |

\* numbering includes 12 aa N-terminal FLAG sequence, n.d. not determined

The replacement of 5 amino acid residues in the course of generating SK2_MOD (see Methods section) from wild type human SphK2 did not impact the assay results. Nearly all the deletion mutants did not grow on galactose (suggesting functional enzyme). Only a single deletion mutant – that lacking both the N-terminal and C-terminal segments (mutant 33-1) – grew on galactose solid media (not shown), which indicates that the 33-1 mutant enzyme has low or absent kinetic activity in yeast. All other mutants except 15-1 were rescued to some extent by the SphK2 inhibitor, SLM6031434, albeit in some cases with diminished potency. For example, in mutant 11-1, 191 aa were deleted (125 aa from the N-terminus, 66 aa from the central segment) and the enzyme was functional and fully rescued by SLM6031434 with about a 3-fold loss of potency. Deletion of the 44 aa C-terminal region (437-481) of the central segment had a particularly severe, negative effect on the potency of the inhibitor (*e.g*. mutants 26-1, 29-1 and 15-1). An observation that we do not understand is the consistently high background of mutant species 20-1 (N-terminal deletion) on galactose liquid media.

**Fig. 8.**
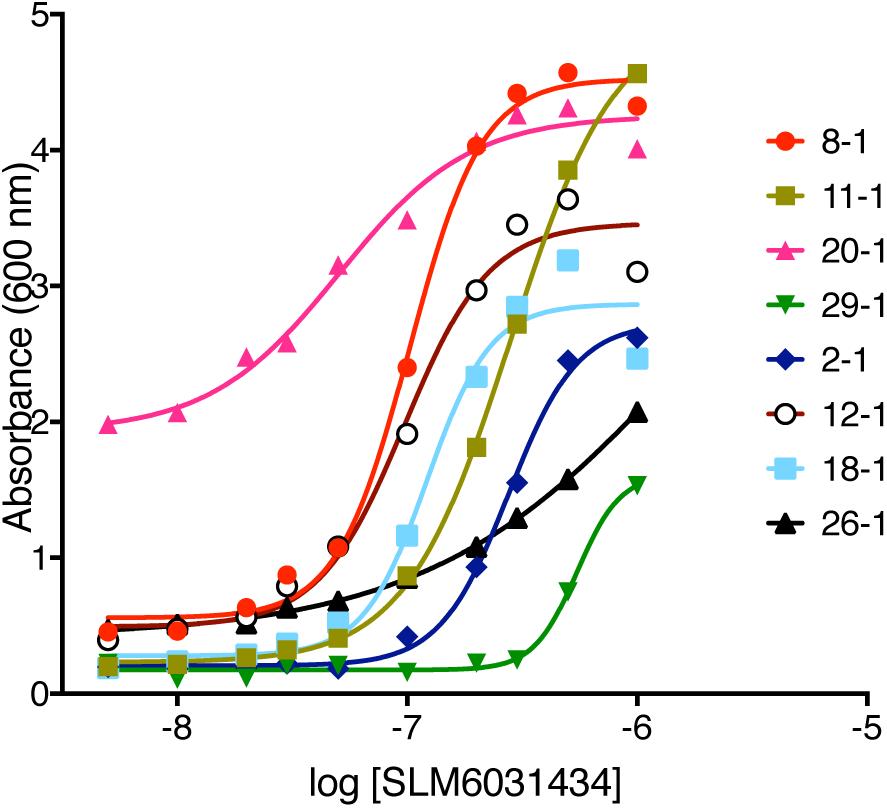
Rescue curves of SK2_MOD deletions mutants by SLM6031434 on the CBY169 strain background.

## Discussion

*Saccharomyces cerevisiae* provides a convenient platform for assessing sphingolipid kinase inhibitors. As predicted by previous studies, expression of sphingosine kinases in mutant strains incapable of degrading phospho-LCBs results in toxic levels that are reduced sufficiently by inhibitors to enable growth. We document herein that ceramide kinases are toxic for a standard laboratory strain of yeast (JS1256) and although we presume this toxicity correlates with the accumulation of phospho-dihydroceramide species, this remains to be proven. The assay is particularly useful for screening, but it can also be used in structure-activity profiling of new chemical entities and for conveniently analyzing mutant sphingolipid kinases.

The yeast-based assay is inexpensive and it requires neither specialized equipment nor radioactive material. Although not rapid (24-48 hours), we have found the assay to be robust, indeed, it yielded reliable results in the hands of four undergraduate research assistants. We were unable to reproduce one minor aspect of the Kashem *et al*. [7] study. That is, we could not achieve sufficient growth of the quadruple null (CBY169) strain seed cultures in media containing raffinose as the fermentable carbon source. We investigated extending the incubation time of the raffinose media seed cultures as well as growing overnight seed cultures in glucose media and switching to raffinose media for two hours before starting the assay. Ultimately, we settled on the expedient strategy of growing overnight seed cultures in glucose media and simply diluting (1:100) these cultures into galactose media to begin our assay.

This maneuver probably is responsible for the high, and somewhat variable background (A_600_ 0.05 – 0.5) in the assay, but there remains a 10-fold differential between minimal and maximal (A_600_ 4-6) responses in the tube assay. Further, we did find consistently that both mouse SphK1 and mouse SphK2 grew slower in galactose media than their human orthologs, which necessitated a 40-48 hour rather than a 24 hour assay time but did not otherwise negatively influence the assay. The assay is readily reduced to a 96 well format and Kashem *et al*. have reported that the assay can be reduced further to a 384 well format using a viability dye in conjunction with a fluorescence based plate reader [7].

The application of the assay suggested initially was to screen chemical libraries for inhibitors of human SphK1 [7]. Our studies suggest that screening for inhibitors of other sphingolipid kinase would also be feasible with this assay – indeed, we have done so with two trypanosomatid sphingosine kinases and human CerK (Kharel, Agah, Barkey-Bircan, Santos and Lynch, unpublished). Screening requires only that compounds accumulate in yeast and not be overtly cytotoxic – success is not strongly dependent on a correlation of inhibitor affinities and the EC_50_ values determined with the assay. Kashem *et al*. [7] suggest that a failure of cytotoxic compounds to be detected is a positive feature of the assay. We agree, but only to the extent that cytotoxicity in yeast predicts cytotoxicity in mammalian cells rather than reflecting some peculiarity of yeast biology, which would be an impediment rather than an advantage.

In contrast to screening chemical libraries, the usefulness of this assay in assessing new chemical entities from a sphingolipid kinase inhibitor medchem campaign depends strongly on the facility of the yeast assay in reporting rank order potencies that reflect the rank order potencies based on inhibitor affinities. In this regard, we are encouraged by our results with one chemical scaffold (1-guanidino-2-phenyloxadiazolylpyrrolidines, *e.g.* SLM6031434) but we were concerned initially about the (*R)*-prolinol scaffold (1-benzyl-2-methanolpyrrolidines, *e.g.* PF-543) because of the large discrepancy between EC_50_ and K_I_ values for PF-543. However, subsequent testing of PF-543 analog compounds suggests that the discrepancy is peculiar to PF-543 rather than a general feature the (*R)*-prolinol chemical scaffold (Hao, Kharel, Lynch and Santos, submitted for publication).

The salient limitations of the yeast assay are those of any cell-based assay. Namely, the assay reports EC_50_ values rather than K_m_, V_max_ or k_cat_ values of inhibitors, test compounds must not be markedly cytotoxic at the concentrations tested and compounds must accumulate to some extent in the yeast. Cytotoxicity is readily discerned by testing compounds for an effect on growth of strains without lipid kinase expression. For example, we found that an early generation dual SphK inhibitor, SKI-II, is toxic for yeast strain CBY169 at concentrations of > 3 μM in our assay. Further, we found that compounds such as SLM6031434, SLC5111312 and SLP7111228, which share a pedant 1-guanidino-2-phenyloxadiazolylpyrrolidine group, are likewise toxic at concentrations > 2-3 μM. In the former case, full growth curves cannot be generated and thus an EC_50_ value for SKI-II cannot be determined with this assay. In contrast, as documented in Fig. 2, 3 and 8, the higher potency guanidino compounds do yield full curves, and thus EC_50_ values are obtained.

Accumulation of a test compound in *S. cerevisiae* can be problematic. If lack of accumulation is due to active extrusion (rather than failure to penetrate and/or metabolism), elimination of candidate transporters, *e.g.* PDR5p, sometimes provides relief. For example, the human SphK1 inhibitor, PF-543, did not rescue growth of human SphK1-expressing yeast unless the *PDR5* gene was deleted (see Fig. 2B). In contrast, the potency of the SphK2 inhibitor, SLM6031434, was not different in the presence (CBY169 strain) or absence (KYA1) of *PDR5* (Fig. 3C), but other 1-guanidino-2-phenyloxadiazolylpyrrolidine compounds were substantially more potent with the KYA1 strain (unpublished data). Since new chemical entities that might be extruded by PDR5p are not recognizable *a priori*, prudence dictates that library screening and assessment of new chemical entities be conducted with *PDR5*Δ strains, *e.g.* KYA1. Our failure to detect activity of ABC294640 at human or mouse SphK2 occurred with both the CBY169 and KYA1 strains, thus we do not know whether this SphK2 inhibitor compound is metabolized rapidly by yeast, fails to penetrate the cell or is extruded by another transporter.

Another limitation is the toxicity of DMSO (dimethylsulfoxide) for *Saccharomyces cerevisiae* at concentrations > 2% (v/v) in SC media. This toxicity could be nettlesome in screening chemical libraries, which are often arrayed in this solvent. In addition, some exogenous lipid kinases might not be expressed in yeast due to proteolysis or their forced expression might be toxic independent of enzyme activity. Protein expression is readily determined using antisera directed against the N-terminal FLAG epitope (see Fig. 7B) and non-enzyme related toxicity can be assessed with catalytically inactive mutant enzymes, but either scenario obviates the assay as a tool to study enzyme inhibitors. We have yet to encounter either of these limitations in our experience to date, which includes the sphingolipid lipid kinases described herein as well as two trypanosomatid sphingosine kinases, pig SphK1 and pig SphK2.

Finally, the usefulness of the yeast platform in interrogating sphingosine and ceramide kinases encouraged us to consider additional applications. One of these is to make advantage of the toxicity of these enzymes to assess mutated forms. We are particularly interested in two questions. Specifically, can a single amino acid change ‘humanize’ mouse SphK1 in terms of inhibitor affinity? And, what is the minimum size of functional SphK2? The former question is important because numerous disease models are available only in the mouse, but there are few potent mouse SphK1 inhibitors. Our results suggest that a single amino change, which is readily accomplished with Crispr/Cas9 technology, can humanize mouse Sphk1 for at least one inhibitor and provides a route for rapidly testing other inhibitors. We used the yeast assay to document also that a ‘mini’ SphK2, which approaches the size of SphK1, is enzymatically competent. The SphK2 deletion series that we describe could prove useful in low resolution mapping of monoclonal antibody epitopes as well as guide expression in *E. coli* for crystallization studies.

## Contributions of Authors

SA developed the yeast assay initially in both the tube and 96 well plate formats, JG, OTE and PBB performed assays, YK generated most of the SphK and CerK plasmids, performed assays and analyzed data, KRL performed assays, served as the project leader and wrote the manuscript, AJM deleted *PDR5* in the CBY169 strain to generate the KYA1 strain and performed assays, WLS provided test compounds and JSS provided facilities and general advice on *Saccharomyces cerevisiae* biology. All authors participated in editing the manuscript.

## Grant support

NIH R01 GM104366 & NIH R01 GM121075 to KRL & WLS, NIH R01 GM075240 to JJS, University of Virginia Summer Research Internship Program fellowships to JG and PBB.

### Acknowledgements

The authors thank Kyle Cunningham, Johns Hopkins University, for his gift of strain CBY169, John Swindle, CompleGen, Seattle, WA and Marek Nagiec, Broad Institute, Boston, MA for their advice and to Cungui Mao, Stony Brook University Cancer Center, Stony Brook, NY for his gift of pYES2-FLAG/URA3 plasmid DNA.

## Declared competing interests of authors

none

## Reagent availability

The KYA1 strain with resident plasmids described are available on request. The pYES2 plasmids with sphingolipid kinase inserts have been deposited with Addgene.

